# Are Pacific Chorus Frogs (*Pseudacris regilla*) Resistant to Tetrodotoxin (TTX)? Characterizing Potential TTX Exposure and Resistance in an Ecological Associate of Pacific Newts (*Taricha*)

**DOI:** 10.1101/2022.01.22.475505

**Authors:** Katherine O. Montana, Valeria Ramírez Castañeda, Rebecca D. Tarvin

## Abstract

Animals that frequently encounter toxins often select for mechanisms of toxin resistance. Both predators that consume toxic prey and organisms in physical contact with a toxin or pollutant in their environment may experience natural selection for resistance. Based on field observations that Pacific Chorus Frogs (*Pseudacris regilla*) sometimes eat and mistakenly amplect tetrodotoxin (TTX)-defended *Taricha* newts, we predicted that *P. regilla* may possess resistance to TTX. We tested this prediction by comparing the amino acid sequences of the molecular target of TTX, the muscle voltage-gated sodium channel gene SCN4A (*Na*_*V*_*1.4*), in populations of *P. regilla* that are sympatric and allopatric with *Taricha*. We identified a single substitution in *Na*_*V*_*1.4* of *P. regilla* in a conserved site near the pore loop where TTX binds. Although the role of this site in TTX resistance has not been functionally assessed, both allopatric and sympatric *P. regilla* had this substitution, suggesting that it may be unrelated to TTX exposure from *Taricha*. Thus, there is no conclusive evidence that *P. regilla* has selected for TTX resistance encoded by amino acid substitutions in this domain. In addition, California occurrence data from the last 50 years indicate that *Taricha* activity peaks in January while the activity of *P. regilla* peaks in April. These relatively distinct activity patterns suggest that *P. regilla* may not be exposed to levels of TTX from *Taricha* that are high enough to select for mutations in the sodium channel. Nevertheless, other unidentified mechanisms of TTX resistance could be present in *P. regilla* and other species that are sympatric with toxic newts.

**Resumen:** Los animales que tienen contacto frecuente con toxinas suelen desarrollar mecanismos de resistencia a las mismas. Tanto los depredadores que consumen presas tóxicas como los organismos en contacto cercano con una toxina o contaminante en su entorno pueden experimentar una presión de selección que los lleva a evolucionar resistencia a toxinas. Basándose en las observaciones de que las ranas coro del Pacífico (*Pseudacris regilla*) a veces comen por error y/o amplexan salamandras del género *Taricha* que poseen tetrodotoxina (TTX), se planteó la hipótesis de que *P. regilla* podría poseer resistencia a la TTX. Esta predicción fue probada comparando las secuencias de aminoácidos en el loop del poro del dominio IV en el gen del canal de sodio voltaje dependiente muscular SCN4A (proteína *Na*_*V*_*1.4*) en poblaciones de *P. regilla* que son simpátricas y alopátricas con *Taricha*. Se identificó una única sustitución en el *Na*_*V*_*1.4* de P. regilla en un sitio conservado cerca del loop del poro donde se une la TTX. Aunque el papel de este sitio en la resistencia a la TTX no ha sido evaluado funcionalmente, tanto el *P. regilla* alopátrico como el simpátrico tienen esta sustitución, lo que sugiere que no está relacionado con la exposición a la TTX secretada por *Taricha*. Por lo tanto, no hay evidencias concluyentes de que *P. regilla* haya evolucionado resistencia a la TTX por medio de sustituciones de aminoácidos en este dominio. Por otro lado, los datos de ocurrencia en California de la actividad de *Taricha* en los últimos 50 años indican alcanza su máximo en enero, mientras que la de *P. regilla* lo hace en abril. Estos patrones de actividad relativamente distintos sugieren que *P. regilla* puede no estar expuesta a niveles de TTX provenientes de *Taricha* que sean lo suficientemente altos como para inducir la evolución de mutaciones en el canal de sodio. Sin embargo, otros mecanismos no identificados de resistencia a la TTX podrían estar presentes en *P. regilla* y en otras especies simpáticas a los salamandras tóxicas.

*Palabras clave: Resistencia a las toxinas; California; Toxinas ambientales; Insensibilidad en el sitio de union; Salamandras; Ecología química*

---

Newts of the genus *Taricha* (*Taricha torosa, T. granulosa, T. sierrae*, and *T. rivularis*, herein collectively referred to as *Taricha*) wield tetrodotoxin (TTX) on their skin (Brodie et al., 1974; Mailho-Fontana et al., 2019; Reimche et al., 2020) that likely functions as an anti-predator defense (Brodie et al., 2002; Williams et al., 2010; Bucciarelli et al., 2017). TTX is a neurotoxin that blocks voltage-gated sodium channels by binding to highly conserved regions of the channel via hydrophobic forces and hydrogen bonds, fitting in the opening of the channel and blocking proper transport of Na^+^ ions (Kao and Levinson, 1986). TTX binding prevents action potentials from propagating, resulting in cessation of nerve signals (Kao and Levinson, 1986). In vertebrate animals lacking TTX resistance, ingested TTX can temporarily paralyze or, in some cases, kill the organism. The poison usually becomes fatal when respiratory muscles fail, resulting in suffocation (Brodie, 1968). The San Francisco Bay Area in California is one of two known hotspots of the arms race between newts and garter snake predators (Brodie et al., 2002; but see Bucciarelli et al. 2022). Here we explore the idea that high levels of TTX defenses in newts may select for resistance in other sympatric organisms outside the predator-prey dynamic, namely the Pacific Chorus Frog, *Pseudacris regilla*.

*Taricha* and their relatives possess amino acid changes in the muscle voltage-gated sodium channel *Na*_*V*_*1.4* that help them resist their own TTX defenses (Hanifin and Gilly, 2015; Gendreau et al., 2021). Amino acid substitutions in voltage-gated sodium channels (VGSCs) alter the shape of the pore loop (p-loop) and the surrounding area which prevents TTX from binding, especially in domain IV (DIV) of *Na*_*V*_*1.4* (Geffeney et al., 2005; Tikhonov and Zhorov, 2005; Jost et. al, 2008; Hanifin and Gilly, 2015). Amino acid substitutions in the p-loop can affect the function of voltage-gated sodium channels by altering sodium ion selectivity or gating properties; hence, trade-offs may exist between TTX resistance and organismal function (Chiamvimonvat et al., 1996; Lee et al., 2011; Hague et al., 2018; but see Moniz et al., 2021). Thus, amino acid changes are often confined to specific conserved regions that can provide resistance while allowing the organism to maintain sodium channel functionality (Feldman et al., 2012). The presence of amino acid substitutions that alter toxin binding is a mechanism known as target-site insensitivity and is the focus of this study.

TTX from *Taricha* skin has been shown to enter waterways and affect the behavior of organisms present in the same aquatic systems. Larval *T. torosa* use TTX as a cue to detect and avoid adult *T. torosa* that sometimes cannibalize larvae (Zimmer et al., 2006). TTX in water can also slow dragonfly nymph consumption of *T. torosa* larvae (Bucciarelli and Kats, 2015). The New Zealand Mud Snail (*Potamopyrgus antipodarum*) moved out of its stream environment when encountering biologically relevant levels of TTX or chemical cues from *T. torosa* containing TTX (Ota et al., 2018). Water containing TTX also increases newt trematode parasite mortality (Calhoun et al., 2017). Thus, naturally occurring levels of TTX produced by *T. torosa* can affect other organisms in the same aquatic environment.

While *P. regilla* do not regularly prey upon *Taricha* (or vice versa), they likely come into contact with TTX in pond water shared with *Taricha* or via direct contact with *Taricha*, sometimes in mistaken amplexus behavior or even attempted predation (Fig. 1). Given that *P. regilla* have permeable skin, we expected that they would be exposed to TTX through physical contact with newts or by occupying pond water that contains newts. We hypothesized that TTX exposure has selected for TTX resistance in *P. regilla* populations that are microsympatric with *Taricha*. We aimed to test whether the sympatry of *Taricha* newts with *Pseudacris regilla* has selected for target-site insensitivity in the domain IV of the skeletal muscle voltage-gated sodium channel (*Na*_*V*_*1.4*) of *P. regilla*.

**Fig. 1.**
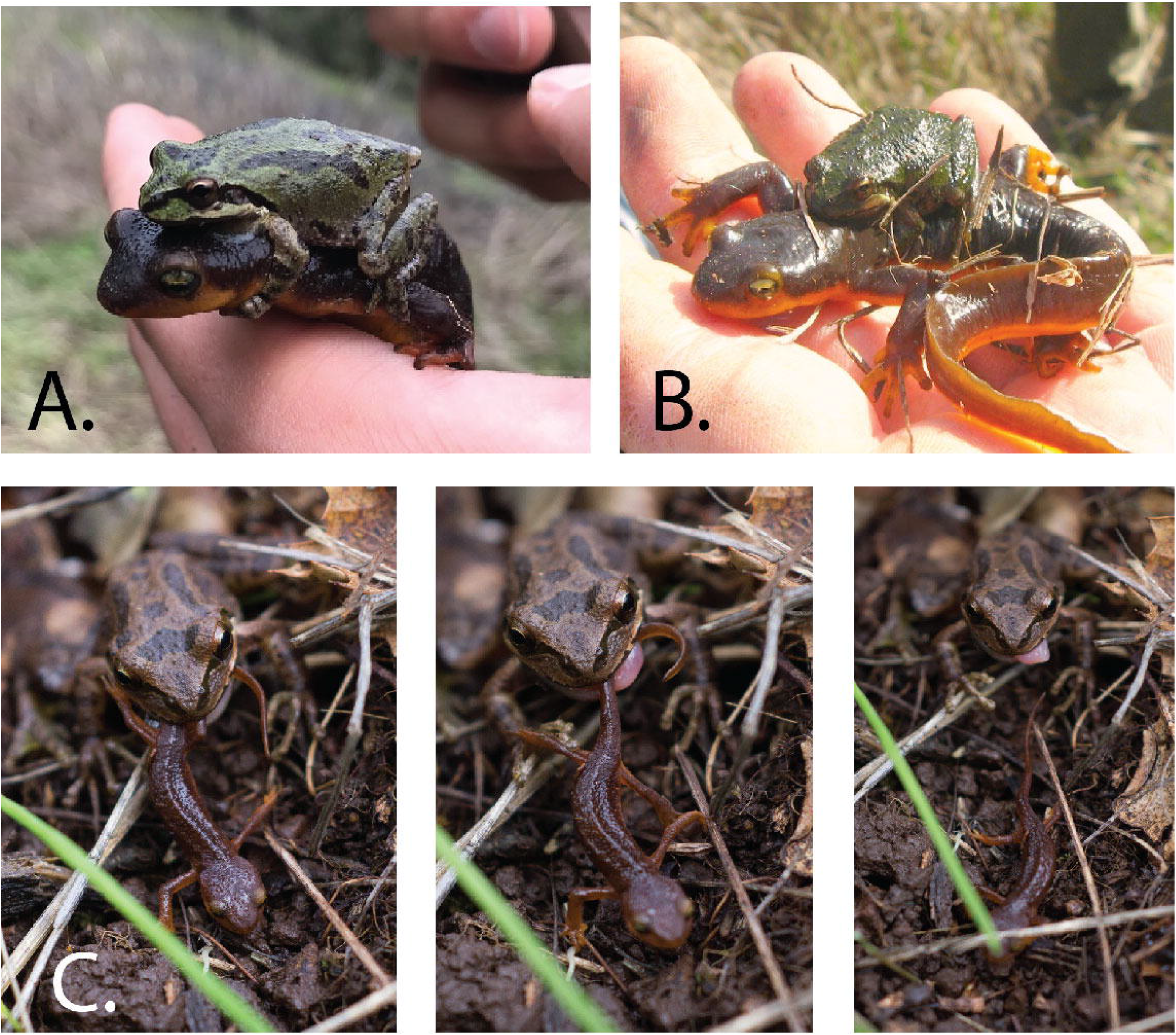
Examples of *Pseudacris regilla* and *Taricha torosa* interactions: A. Amplexus in Briones Regional Park, Martinez, CA, 1 March 2019, photo credit K. Montana, user kmontana on iNaturalist (CC BY-NC 4.0; observation 100725148); B. Another instance of amplexus in Briones Regional Park, Martinez, CA, 21 February 2009, photo credit user ap2il on iNaturalist (CC BY-NC 4.0; observation 550227); C. Attempted predation on *Taricha torosa* by *Pseudacris regilla* in Mark West Springs, CA, 1 March 2014, photo credit user tiwane on iNaturalist (CC BY-NC 4.0; observation 2312). The frogs did not appear to be harmed in any of these scenarios, though the tongue posture in C suggests some numbing effects from newt exposure.

Our study centered on California populations, examining *Taricha* and *P. regilla* that do and do not overlap geographically in the present. We conducted demographic surveys of *Taricha granulosa, T. torosa*, and *P. regilla* at three San Francisco Bay Area ponds where the species co-occur to estimate the extent to which *P. regilla* may encounter TTX. We supplemented our surveys with occurrence data available through the Global Biodiversity Information Facility (GBIF) to better estimate overlap of annual activity patterns and to determine the degree of sympatry between *Taricha* spp. and *P. regilla*. To determine whether *P. regilla* has structural modifications in the muscle voltage-gated sodium channel *Na*_*V*_*1.4* that may provide TTX resistance, we sequenced and reviewed amino acid residues encoded by the SCN4A gene. Some amino acid substitutions in domain IV of SCN4A are known to confer resistance to TTX and have been found in many organisms that either have TTX defense or consume TTX-defended prey (Geffeney et al., 2005; Venkatesh et al., 2005; Jost et al., 2008; Feldman et al., 2012; Hanifin and Gilly, 2015; Gendreau et al., 2021). We hypothesized that if the presence of *Taricha* selects for TTX resistance in *P. regilla*, populations sympatric with *Taricha* would have mutations in the domain IV p-loop of *Na*_*V*_*1.4* while populations that are allopatric would not. Our work provides insight into the potential cascading ecological consequences and unintended effects of chemical defenses.

## Materials and Methods

### Field Work

We studied *Taricha* and *P. regilla* at four San Francisco Bay Area ponds. We conducted surveys at three of these sites where *Taricha granulosa, T. torosa*, and *P. regilla* co-occur: two sites in Briones Regional Park in Martinez, Contra Costa County, CA (Old Briones Road Trail 1 [OBRT 1] at 37.94571, -122.13395, and OBRT 2 at 37.94352, -122.14102, named after the trail closest to the sites) and one site at the Japanese Pool (37.87440, -122.23760) at the University of California Botanical Garden (UCBG) in Berkeley, Alameda County, CA (Fig. S1). OBRT 1 and OBRT 2 are in an unshaded area of grazed open grassland that lies within an oak woodland habitat, and they connect to each other when water levels are high. The UCBG Japanese Pool is a human-constructed pool with a concrete floor and sides, located under tree cover within the visitor area of the UCBG (Fig. S1). We selected these three ponds because we knew them to contain populations of *P. regilla* as well as *Taricha torosa* and *Taricha granulosa*. Close to our field sites, *P. regilla* and *T. torosa* were known to be prey of *Thamnophis* (Preston and Johnson, 2012), supporting cohabitation in the San Francisco Bay Area. We also collected specimens from a fourth pond (Mountain Lake) located in the Presidio of San Francisco (37.788496, -122.468605) that has historically contained *P. regilla* but no *Taricha* (records show absence of *Taricha* since at least 1966; pers. comm. J. Young, Presidio Trust). As of 2021 (shortly after our surveys), efforts were being made to reintroduce *Taricha* to the Presidio waterways. Mountain Lake is a natural lake surrounded by grass and receives run-off from the nearby Presidio Golf Course (Fig. S1). From October 2019 to February 2020, we conducted six surveys at the two Briones ponds and five surveys at UCBG, with surveys at each site occurring three to four weeks apart from one another. We were unable to gather abundance data for more than this time period because the parks were closed from February until October 2020 during the early months of the COVID-19 pandemic.

Two researchers (usually K. Montana and V. Ramírez-Castañeda) conducted surveys in the morning or early afternoon (between 8:00 AM and 1:30 PM). During each survey, surveyors split the pond perimeter into 7.62-m (25-feet) sections, and circled the pond on foot, counting how many individuals of each species were visible in the water from the shore within each transect. We note that surveyors were only able to count the number of visible newts. OBRT 1 and OBRT 2 were murky with plants and algae in which newts tended to hide. Although the surface of the Japanese Pool at UCBG was glassy, the pool was filled with plants that likely obscured observations. Surveyors also counted the numbers of visible, individual adult *Pseudacris regilla*; *P. regilla* that were calling but not visible were not counted because surveyors could not ascertain if multiple calls were coming from a single or a few individuals. For the purposes of this study, we did not differentiate between *T. torosa* and *T. granulosa* when we recorded their occurrences because they both contribute to the presence of TTX in pond water, and we could not easily identify individuals to species from the pond shores. We counted large (∼5 cm or longer) *Taricha* larvae, adult *Taricha*, and *Taricha torosa* egg masses, which are conspicuous groups of eggs known to possess TTX via vertical transfer from the parent (Gall et al., 2012). The much smaller *T. granulosa* eggs are typically hidden in the substrate and are more difficult to see; hence we did not include them in our survey. To simplify our survey protocol, we counted larval *Taricha*, which also have some amount of TTX (Gall et al., 2011), together with adults. We plotted survey results using R version 4.1.1 and the packages ggplot2 v.3.3.5 and ggpubr v.0.4.0 (Wickham, 2016; Kassambara, 2020; R Core Team, 2021) (Fig. S2).

After COVID-19 conditions permitted field work to continue in October 2020, we collected one *P. regilla* from the dry bed of OBRT 2 and eight *P. regilla* from the east arm of Mountain Lake in the Presidio of San Francisco. We surveyed OBRT 1 and UCBG at the same time but did not find any *P. regilla* at either location; OBRT 1 was dry.

### GBIF Occurrences and Sympatry

We estimated seasonal patterns of *P. regilla* and *Taricha* activity across the state of California and within the San Francisco Bay Area (N 38.66226, S 37.09298, E -121.12701, W -123.23579) by reviewing annual patterns of GBIF occurrence data of *P. regilla* and *Taricha* from 1 January 1971 to 23 November 2021 (GBIF) using the following R packages: RColorBrewer v.1.1.2, ggplot2 v.3.3.5, chron v.2.3.56, cowplot v.1.1.1, and dplyr v.1.0.7 (Neuwirth, 2014; Wickham, 2016; James and Hornik, 2020; Wilke, 2020; Wickham et al., 2021). In our search terms for GBIF, we took into account that the taxonomy of *Pseudacris regilla* has changed several times over the last few decades (Recuero et al., 2006; Fouquette and Dubois, 2014; Duellman et al., 2016). iNaturalist categorizes some *Pseudacris regilla* (e.g., in the San Francisco Bay Area) as *Pseudacris sierra*. However, these distinctions are not universally accepted by the herpetology community (Barrow et al., 2014; Crother et al., 2017). Here we consider *P. sierra, P. hypochondriaca*, and *Hyla regilla* synonyms of *Pseudacris regilla*. We downloaded occurrence data from GBIF for the *P. regilla* synonyms listed here and for all currently recognized species of *Taricha* (*T. granulosa, T. rivularis, T. sierrae, T. torosa*), as well as those no longer recognized (*T. lindoei, T. miocenica*, and *Palaeotaricha oligocenica*) and individuals identified only to the genus *Taricha*. GBIF data include Research Grade iNaturalist observations in addition to museum and specimen records.

We aimed to sequence *Na*_*V*_*1.4* from *P. regilla* that were sympatric and allopatric with *Taricha* (see below). We originally classified *P. regilla* tissue samples as sympatric if they were collected within 35 km or 500 m of elevation from the nearest *Taricha* occurrence documented on GBIF as of 23 November 2021; this included 11 individuals and the single *P. regilla* collected at OBRT 2. We classified allopatric *P. regilla* samples as those individuals collected at least 35 km or 500 m in elevation away from the nearest *Taricha* observation; this included 18 individuals and the eight *P. regilla* from the Mountain Lake field site in the Presidio (Table S1).

To review general patterns in geographic range overlap between species, we plotted *P. regilla* and *Taricha* GBIF occurrence data from the western USA, as well as localities of museum samples we sequenced using ggplot2 v.3.3.5 and tidyverse v.1.3.1 (Wickham, 2016; Wickham et al., 2019; GBIF). We plotted the occurrence data of these species to confirm that the museum tissues were from locations where *P. regilla* and *Taricha* are allopatric or sympatric. Specimens that were collected from localities within overlapping hexagons were considered to be sympatric; none of our categorizations changed. The binwidth of each hexagon was 0.05 in the vertical and horizontal directions. We note that this visualization, though qualitatively useful, is limited in several ways. For example, our designations of sympatry versus allopatry are not indicative of species distributions prior to 1971. Additionally, iNaturalist data included in the GBIF data can be biased; for instance, individuals are unlikely to post multiple images of a single species at a site and observations can be more highly concentrated in regions where the iNaturalist is well-known (Hochmair et al., 2020). Future studies may consider estimating more accurate species distribution models.

### DNA Sequencing and Analysis

We selected 29 liver tissues of *Pseudacris regilla* from the Museum of Vertebrate Zoology at the University of California, Berkeley to broaden our geographic sampling beyond the San Francisco Bay Area (Table S1). In total, we sequenced *Na*_*V*_*1.4* from 11 *P. regilla* that were sympatric with *Taricha* and 18 that were allopatric. We predicted that *Na*_*V*_*1.4* sequences from sympatric, but not allopatric, *P. regilla* would show amino acid substitutions potentially conferring TTX resistance. We also sequenced *Na*_*V*_*1.4* from two close relatives of *P. regilla, Pseudacris cadaverina* and *Acris crepitans* (Table S1). The individuals of *P. cadaverina* and *A. crepitans* are from locations far from *Taricha* and, thus, were likely not exposed to TTX from *Taricha*. However, the range of *A. crepitans* overlaps with another TTX-bearing newt, *Notophthalmus viridescens* (Baird, 1854; Mecham, 1967), so it may be exposed to TTX as well. Thus, if TTX from newts selects for mutations in *Na*_*V*_*1.4* in sympatric frog species, we would expect to find amino acid substitutions in *Na*_*V*_*1.4* of *A. crepitans* as well as in *Na*_*V*_*1.4* of *P. regilla* sympatric with *Taricha*. In contrast, we expected to find no resistance-conferring amino acid substitutions in allopatric *P. regilla* or in *P. cadaverina*.

In the Evolutionary Genetics Lab (UC Berkeley), we extracted DNA from tissues according to the Miller et al. (1988) salt-extraction protocols, with modifications (see Supporting Methods for details). The primers used for *P. regilla* samples targeted exon 24 (DIV segment 6 and p-loop) of the gene SCN4A: 2268F_SCN4A 5’-TCTCCCGGCCCTCTTCAATA-3’ and 2681R_SCN4A 5’-TCGTCCTCGCATAAAGGCTC-3’ (Tarvin et al., 2016). We added 0.5 µL of MgCl_2_ to each PCR sample. PCR protocols were: an initial denaturation for 2 min at 94 ºC; 35 cycles of: 94 ºC at 30 s for denaturation, an annealing temperature of 54 ºC for 30 s, and an elongation temperature of 72 ºC for 45 s; and a final extension time of 7 min at 72 ºC. PCR products were run on an agarose gel, purified with ExoSAP-IT PCR Product Cleanup Reagent (ThermoFisher Scientific, 78201.1.ML) and sequenced using the reverse primer 2681R_SCN4A at the UC Berkeley DNA Sequencing Facility. For PCR and sequencing samples of *Pseudacris cadaverina* and *Acris crepitans*, we used the same protocol and forward primer, but the reverse primer was redesigned based on the DIV SCN4A *P. regilla* sequences we produced: P.regilla_SCN4AD4_R 5’-ACTGCTTTCTTCTGTGGCCAC-3’.

Using BLAST (Altschul et al., 1990) with the *P. regilla Na*_*V*_*1.4* sequence as a query, we obtained 180 previously published representative sequences of SCN4A from other herpetofauna including salamandrids, dendrobatids, snakes, and lizards with available data on GenBank and DataDryad (Table S2; Gendreau et al., 2021; Gendreau et al., 2021). We compare these SCN4A sequences, as well as those of *P. cadaverina* and *A. crepitans*, to SCN4A of our *P. regilla* samples. Of the additional 180 sequences added, *Erythrolamprus epinephelus, Rhabdophis tigrinus, Thamnophis elegans, Thamnophis sirtalis, Thamnophis atratus, Taricha torosa, Taricha granulosa, S. salamandra, Tylototriton shanjing*, and other salamandrids are either confirmed or hypothesized to possess target-site insensitivity to TTX via mutations in *Na*_*V*_*1.4* (Feldman et al., 2009; Feldman et al., 2012; Hanifin and Gilly, 2015; Ramírez-Castañeda, 2017; Gendreau et al., 2021; Vaelli et al. 2021). We use *Rattus norvegicus* as a reference (Uniprot #P15390) of a species that has not been documented to be resistant to TTX.

We identified the location of the DIV p-loop in *Na*_*V*_*1.4* based on protein domains inferred by Tikhonov and Zhorov (2005), and we numbered the residues according to SCN4A of Rattus norvegicus (UniProt accession #P15390). We aligned sequences using Clustal Omega in Geneious (Sievers et al., 2011, Geneious Prime 2020.1.1). We reviewed the alignment by eye for amino acid differences between sequences, focusing on residues that have been shown to provide TTX resistance. We estimated a phylogeny of the *Na*_*V*_*1.4* DIV sequences using maximum likelihood on the IQTree web server, with two partitions (one for codon positions 1 and 2, another for codon position 3), the GTR + G substitution model, and support assessed using 1000 ultrafast bootstraps and the SH-aLRT test (Nguyen et al., 2015; Chernomor et al., 2015; Trifinopoulos et al., 2016; Hoang et al., 2018; IQ-Tree web server http://iqtree.cibiv.univie.ac.at/). We edited and visualized the phylogeny using the Interactive Tree of Life (Letunic and Bork, 2021).

## Results

### Field Surveys

While we did not observe *P. regilla* at OBRT 2 or UCBG, their abundance at OBRT 1 peaked early in the observation period at 32 individuals on the first field trip date (20 October 2019). For OBRT field sites, *Taricha* abundance peaked at the beginning of February; for the UCBG, abundance was low, and we may not have observed the seasonal peak (surveys were suspended at the end of February 2020 because of the COVID-19 pandemic). *Taricha torosa* egg masses were most abundant at the beginning of February 2020 at OBRT 1 and OBRT 2, while they were most abundant at UCBG at the end of February (though we cannot be certain these are true peaks because our surveys ended before March) (Fig. S2).

### GBIF

California-wide GBIF observations for *P. regilla* and *Taricha* from 1 January 1971 to 23 November 2021 indicate two seasonal peaks of *P. regilla* abundance in April and August, while *Taricha* peaks in January (Fig. S3). Though the abundance peaks do not occur at precisely the same time, *P. regilla* and *Taricha* do overlap with one another and, thus, the species have opportunities to interact such that *Taricha* could expose *P. regilla* to TTX. These data are summarized across California and thus represent average activity across a range of habitat types. Nevertheless, when the data are constrained to a square roughly representing the San Francisco Bay Area (N 38.66226, S 37.09298, E -121.12701, W -123.23579), the results are similar, indicating a peak for *Taricha* in January and a peak for *P. regilla* in March (Fig. S4). Along the west coast of the USA, patterns of geographic sympatry estimated from GBIF occurrence data vary substantially (Fig. 2). Hot spots of sympatry were found in Humboldt County, the San Francisco Bay Area, Los Angeles area, and northern Oregon, although much of each taxon’s range does not overlap with that of the other.

**Fig. 2.**
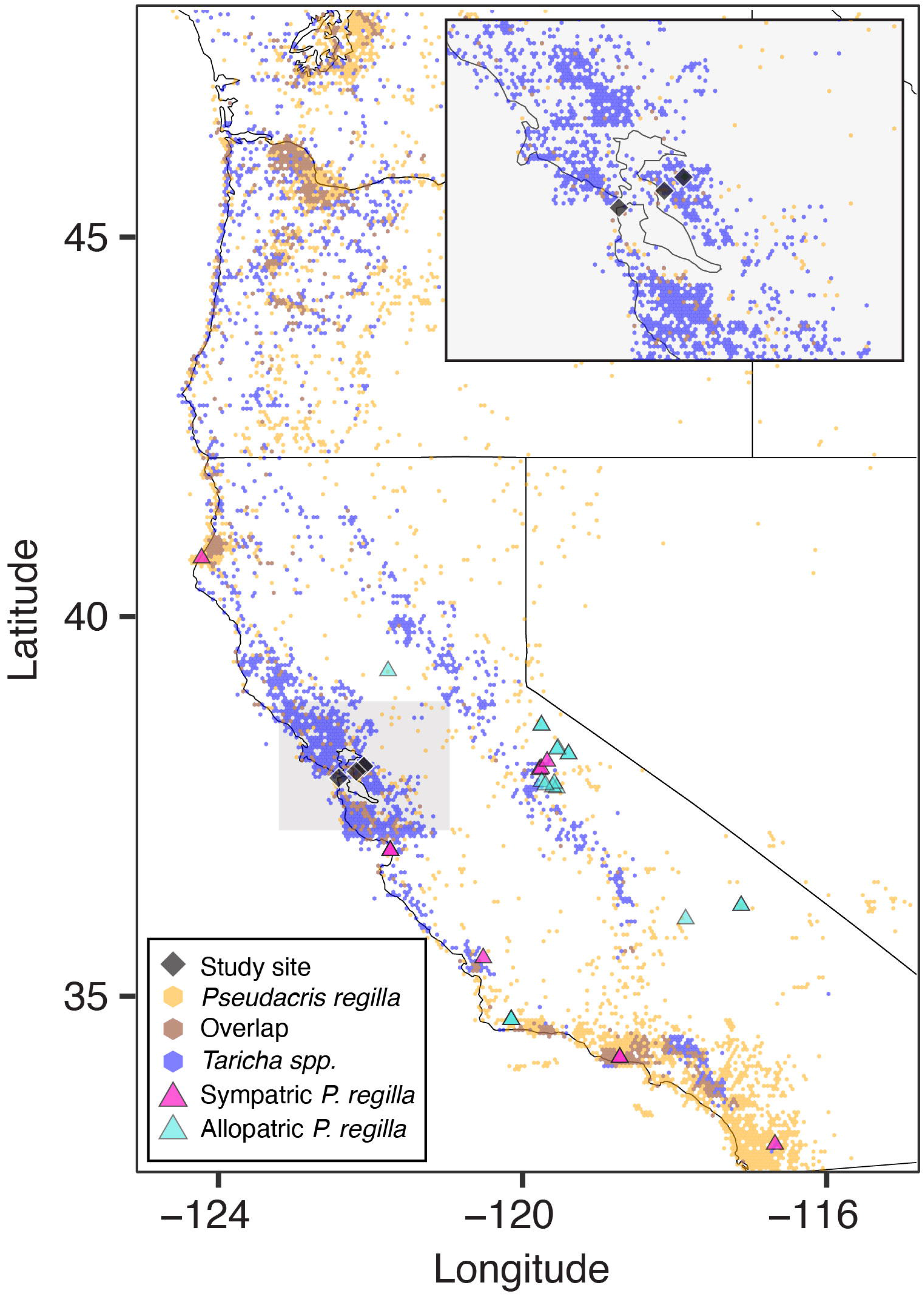
Map showing overlap of *P. regilla* and *Taricha* occurrences on the west coast of the USA. Data are from GBIF, collected through 23 May 2022 for *Taricha* and through 6 June 2022 for *P. regilla. P. regilla* museum specimens that are sympatric with *Taricha* are indicated by pink triangles; *P. regilla* museum specimens allopatric with *Taricha* are indicated by blue triangles; field sites are indicated by black squares. Geographic regions with only *Taricha* occurrences are indicated by blue hexagons; those with only *P. regilla* are indicated by orange hexagons; regions where *Taricha* and *P. regilla* overlap are indicated by brown hexagons.

### Amino Acid Changes

We found one amino acid change (I1519M, numbered according to the *Rattus norvegicus* UniProt accession #P15390) within the DIV p-loop of the sodium channel protein *Na*_*V*_*1.4* in *Pseudacris regilla* compared to a sequence of *Rattus norvegicus* (a common species used in comparative TTX studies [e.g., Tikhonov and Zhorov, 2005; Tarvin et al. 2016] as it represents the mammalian TTX-sensitive *Na*_*V*_*1.4* [Chahine et al., 1994]) (Fig. 3). Importantly, there were no observed differences in amino acid sequences between *P. regilla* specimens that were sympatric and allopatric with *Taricha*. Close relatives of *P. regilla* (*P. cadaverina* or *A. crepitans*) did not possess the I1519M substitution, suggesting that the mutation arose within *P. regilla* or its ancestor.

**Fig. 3.**
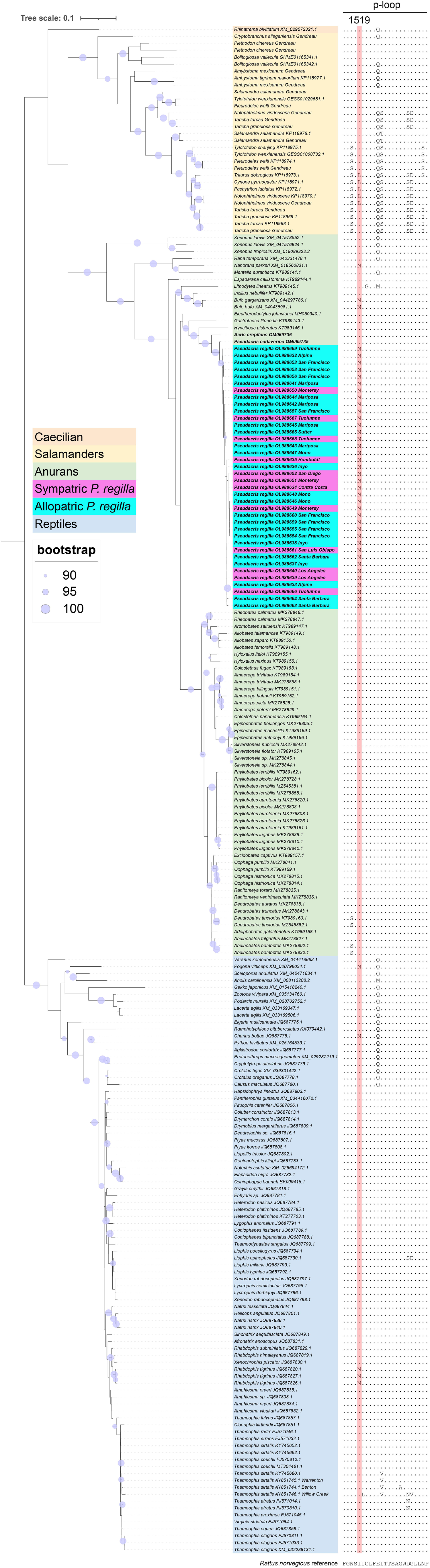
Alignment of the p-loop of domain IV of the voltage-gated sodium channel *Na*_*V*_*1.4* in several reptile and amphibian species including new sequences from *Pseudacris regilla*, mapped onto a phylogeny inferred using 381 bp of the domain IV in *Na*_*V*_*1.4* and IQTREE software (support values are SH-aLRT/ultrafast bootstrap values from 1000 iterations each). Amino acid positions are numbered according to *Rattus norvegicus*, UniProt accession #P15390. The amino acid substitution I1519M is found in *P. regilla* as well as *N. parkeri, Bufo gargarizans, Bufo bufo, Pogona vitticeps, Charina bottae*, and *R. tigrinus*.

## Discussion

The *Taricha-Thamnophis* system is a well-studied example of predator prey interactions involving a toxin. Here we extend the system to include an ecological associate, *P. regilla*. We note that *Taricha* are well known to vary the level of their toxic defenses over time, complicating the arms race story (Bucciarelli et al., 2021; Bucciarelli et al., 2022). Furthermore, while TTX is known to be produced by bacterial symbionts in several marine organisms and in *Taricha torosa* (Magarlamov et al., 2017; Vaelli et al., 2020; Bucciarelli et al., 2021), the biosynthesis pathways of TTX remain unresolved, and endogenous production of TTX in newts is also plausible (Kudo et al., 2020; Kudo et al., 2021; but see Bucciarelli et al. 2022). Here we document ecologically relevant interactions between *Taricha* and *P. regilla* (Fig. 1) with new field observations and an assessment of GBIF data (Fig. 2; Figs. S2–S4). We show the absence of any obvious TTX-resistance-conferring amino acid mutations in the *Na*_*V*_*1.4* DIV p-loop of *P. regilla* (Fig. 3). It remains unknown how *P. regilla* survives living in such close proximity with toxic *Taricha* newts and whether it encounters high enough levels of TTX to evolve resistance. Our data have added to the complex story involving newt toxicity and its effects on members of the surrounding community.

Abundance data indicate that *P. regilla* and *Taricha* annual activity patterns overlap but that peak breeding times are not synchronized (Figs. S2–S4). When we surveyed the field sites in Briones Regional Park and the UC Botanical Garden from October 2019 to March 2020, California was experiencing drought. The area of the ponds fluctuated due to changes in precipitation and, in the case of the Japanese Pool, some drainage for maintenance (UCBG personnel, pers. com.). Thus, weather and pond conditions may have influenced our survey observations.

Observations of mistaken amplexus and predation of newts by *P. regilla* indicate that the species do come into contact (Fig. 1). Still, TTX exposure may not be frequent or potent enough for *P. regilla* to undergo strong selection for resistance to TTX. In cases where environmental selective regimes are not predictable, more plastic responses like upregulation of detoxifying enzymes may be the primary way animals deal with toxin exposure (Zanger and Schwab, 2013). Nevertheless, even occasional exposure to TTX can result in maintenance of target-site insensitivity (Durso et al., 2021).

We hypothesized that we would find TTX-resistance-conferring amino acid substitutions in *Na*_*V*_*1.4* in *P. regilla* that are sympatric with *Taricha* and no substitutions in *P. regilla* that are allopatric with *Taricha*, with a greater number of substitutions indicating a higher level of resistance as has been shown various Na_V_ channels in salamanders and snakes (Geffeney et al., 2005; Feldman et al., 2010; Hanifin and Gilly, 2015; McGlothlin et al., 2016). Compared to the TTX-sensitive *Rattus norvegicus Na*_*V*_*1.4* DIV, we found one amino acid change, I1519M, at a conserved hydrophobic residue located in proximity to TTX-sensing residues in the p-loop (Tikhonov and Zhorov, 2005; Geffeney et al., 2005; Tikhonov and Zhorov, 2011; Feldman et al. 2012; Gendreau et al., 2021). This substitution was present in *P. regilla* that were sympatric and allopatric with *Taricha*. Amino acid changes in important regions of the *Na*_*V*_*1.4* protein do not always translate to a resistant phenotype (e.g., see Abderemane-Ali et al., 2021). Furthermore, the presence of this substitution in all *P. regilla* samples does not support our hypothesis that substitutions would only be found where *P. regilla* is exposed to TTX from sympatric *Taricha* (at least based on occurrence data from the last 50 years). We recognize that our hypothesis made a simplifying assumption that the designation of *P. regilla* populations as sympatric or allopatric with *Taricha* is relevant and valid during the time in which selection may have occurred for TTX resistance, and that gene flow does not occur between sympatric and allopatric *P. regilla* populations. Nonetheless, our study suggests that the I1519M substitution evolved in the ancestor of *P. regilla*, which likely rules out contemporary selection from *Taricha*. Furthermore, our survey of SCN4A sequences in reptiles and amphibians (Fig. 3) suggests that this site is relatively less conserved than other adjacent sites.

In addition to *P. regilla*, the I1519M substitution is present in *Nanorana parkeri, Bufo bufo, Bufo gargarizans, Pogona vitticeps, Charina bottae*, and *Rhabdophis tigrinus* (Fig. 3). None of these species are known for having a strong association with TTX; though, *R. tigrinus* has been observed consuming TTX-defended treefrogs (Feldman et al., 2012). Furthermore, *Nanorana parkeri*, which also possesses the I1519M transition, occurs within the range of the toxic and relatively TTX-resistant newt *Tylototriton verrucosus*, though its toxin is yet to be identified (Brodie et al., 1984; Hanifin and Gilly, 2015, which studies a relative *T. shanjing*). Similarly, *A. crepitans* co-occurs with the TTX-defended newt *Notophthalmus viridescens* (Mebs et al., 2010), but this species does not possess the I1519M substitution.

The close proximity of the I1519M substitution to other changes implicated in TTX resistance may support its importance in TTX resistance (Feldman et al. 2012). However, the hypothesis that a I1519M substitution contributes to TTX resistance has been suggested to be incorrect because the same substitution is found in other TTX-sensitive voltage-gated sodium channel paralogs (McGlothlin et al., 2016; noted as I1555M in their study). Mutagenesis studies suggest that substituting methionine for isoleucine has limited effect on a protein (Ohmura et al., 2001). Another hydrophobic change in the same position (I1519L) is found in several genera of TTX-resistant newts (Fig. 3: *Triturus dobrogicus, Cynops pyrrhogaster, Pachytriton labiatus, Notophthalmus viridescens*); however, it is hypothesized that this change is not directly related to TTX resistance either (Hanifin and Gilly, 2015). Because there is disagreement whether the I1519M mutation (or the site itself) provides resistance, we cannot say with certainty whether it provides TTX resistance in *P. regilla* without further experimental evidence.

While there is not strong evidence that *P. regilla* possesses target-site insensitivity in the p-loop of DIV in *Na*_*V*_*1.4*, it is plausible that *P. regilla* may have other ways of avoiding the effects of TTX. Other regions of *Na*_*V*_*1.4*, such as domain III, and other voltage-gated sodium channels, such as *Na*_*V*_*1.7*, also harbor amino acid substitutions implicated in TTX resistance (Feldman et al., 2012; McGlothlin et al., 2016). *Pseudacris regilla* could possess the ability to smell TTX, as larval *Taricha* can, in order to avoid it (Zimmer et al., 2006). Alternatively, they could possess a diffusion barrier in the skin or the gut that would prevent high levels of TTX exposure, as occurs in mantids (Mebs et al., 2016). It is also possible that *P. regilla* possesses a binding protein that can scavenge TTX. Other frogs have saxiphillin, a protein that binds saxitoxin, which is structurally similar to TTX. However, saxiphilin is incapable of binding TTX, at least in the animals that have been tested (Mahar et al., 1991; Llewellyn et al., 1997; Abderemane-Ali et al., 2021). Similarly, little is known about the potential interactions between *Acris crepitans* and *Notophthalmus viridescens*, or between *Nanorana parkeri* and *Tylototriton verrucosus*, which could also select for toxin resistance in the frogs. To date, none of these alternative resistance mechanisms have been studied in *Pseudacris, Acris, Nanorana*, or salamandrids.

A number of reasons could explain why *P. regilla* would lack target-site insensitivity in *Na*_*V*_*1.4*. Evolving TTX resistance could be costly to *P. regilla* due to possible alterations in the selectivity filter of sodium channels that frequently occurs when amino acid substitutions are present (Chiamvimonvat et al., 1996; Lee et al., 2011; Feldman et al., 2012; Hague et al., 2018; but see Moniz et al., 2021 for evidence against potential impact of mutations on metabolism of the organism). Evolving costly mutations may require a stronger selective pressure than is currently occurring. Given the relative quantities of TTX on newt skin (up to 5.48 mg/newt [Reimche et al., 2020] or even one recording of 8.551 mg/newt [Stokes, 2015]) compared to TTX levels in newt-containing water (∼1×10^−7^ mol/L [Zimmer et al., 2006]), the environmental toxicity of TTX is likely to cause a weaker selection for TTX resistance than exposure via chemical defenses from prey. The cost of exhibiting TTX resistance combined with the low potential for *P. regilla* to interact with high concentrations of TTX in its environment may explain why we did not unequivocally identify target-site insensitivity in *P. regilla* DIV *Na*_*V*_*1.4*. Future work exploring the effects of TTX as an environmental toxin could include measuring TTX concentrations in newt-containing ponds, performing whole organism or *in vitro* assays of *P. regilla* to determine TTX sensitivity, and testing for other modes of TTX resistance.

## Conclusions

We do not find strong evidence for target-site insensitivity in DIV of the skeletal muscle voltage-gated sodium channel *Na*_*V*_*1.4* of *Pseudacris regilla*. Although it is possible that the I1519M substitution in the p-loop of *Na*_*V*_*1.4* may be related to TTX resistance, *P. regilla* may not encounter TTX from *Taricha* with high enough frequency or magnitude to select for TTX resistance. Future seasonal measurements of pond-water concentrations will help evaluate this hypothesis. If *P. regilla* are indeed exposed to significant levels of TTX, they may have behavioral or physiological mechanisms to survive TTX exposure other than target-site insensitivity in *Na*_*V*_*1.4* DIV. Investigating the environmental toxicity of TTX in the relationship between *P. regilla* and *Taricha* is an opportunity to explore the nexus of ecological interactions, neurotoxins, and resistance mechanisms. The likely absence of target-site insensitivity in *P. regilla* indicates that exposure to tetrodotoxin outside of trophic interactions may not strongly select for fixed mechanisms of toxin resistance.

## Acknowledgments

We affirm the sovereignty of the Karkin, Me-Wuk (Bay Miwok), Muwekma, Chochenyo, Ramaytush, and Ohlone people and the Confederated Villages of Lisjan; we thank them for stewarding the land where we worked and, thus, serving an integral role in the production of this study. We thank L. Smith in the Evolutionary Genetics Lab, as well as C. Spencer and the support of the Museum of Vertebrate Zoology collections staff. We also thank those who supported our work at field sites: UC Botanical Garden staff, including H. Forbes, J. Fong, E. Hupperts, and C. Loughran; East Bay Regional Parks District staff, including T. Lin and J. Miller; and J. Young with the Presidio Trust. We thank the other members of the Tarvin Lab for their editing prowess and general support. We thank A. Shabel for his advice in the early stages of this project, and K. Sanko, J. Toman, and L. Galindo who helped as field assistants. We appreciate the newts and frogs who were the focus of this project and who were sacrificed to offer the knowledge they hold. K. Montana thanks her family and ancestors for their guidance. R. Tarvin was supported by UC Berkeley start-up funds.

The East Bay Regional Parks District authorized the surveys at Briones under permit #19-1044, and staff at the UCBG gave written consent for surveys conducted on the premises. All animals were collected under approved collection permits and protocols: for the National Parks Service Scientific Research and Collecting Permit 2 number PRSF-2020-SCI-0002 within study PRSF-00035, with a written extension after expiration of the original permit, University of California IACUC AUP-2019-08-12457, and California Department of Fish and Wildlife Scientific Collection Permit S-190980001-19111-001.

The Spanish translation was done using www.DeepL.com/Translator (free version), which was then edited by Valeria Ramírez-Castañeda.

## Supplementary Online Material

### Supporting Methods

DNA Extraction Procedures, Adapted from Miller et al., 1988.

*Table S1.—*Metadata for tissues obtained from the Museum of Vertebrate Zoology (UC Berkeley) and from newly collected specimens (indicated by asterisks) that were used to sequence SCN4A exon 24 in this study.

*Table S2.—*Sequences of *Na*_*V*_*1.4* (SCN4A exon 24) obtained from GenBank used in Fig. 3.

*Figure S1.—*Field Sites: A. Old Briones Road Trail 1 (OBRT 1) pond, 37.94441, - 122.13377; B. Old Briones Road Trail 2 (OBRT 2) pond, 37.94352, -122.14102; C. Japanese Pool at the UC Botanical Garden (UCBG), 37.87440, -122.23760; D. Presidio site, 37.788496, -122.468605; all photos taken by K. Montana.

*Figure S2.—*Numbers of A. *P. regilla* adults, B. *Taricha torosa* and *T. granulosa* adults and larvae, and C. *T. torosa* egg masses observed over field trips to the Old Briones Road Trail 1 (OBRT 1) (blue circle line), OBRT 2 (red triangle line), and U.C. Botanical Garden (yellow square line) ponds; 20 October 2019 to late February 2020.

*Figure S3*.—Percentage of total abundance of GBIF-recorded individuals of *Pseudacris regilla* (green) and *Taricha* (blue) in California by month, over the course of the last 50 years. Peak abundances for *P. regilla* occur around April and August each year, and peak abundances for *Taricha* occur in January each year. Samples included were from 1 January 1971 to 23 November 2021.

*Figure S4*.—Percentage of total abundance of GBIF-recorded individuals of *Pseudacris regilla* (green) and *Taricha* (blue) in the San Francisco Bay Area (N 38.66226, S 37.09298, E -121.12701, W -123.23579) by month, over the course of the last 50 years. Peak abundance for *P. regilla* occurs around March each year, and peak abundance for *Taricha* occurs in January each year. Samples included were from 1 January 1971 to 23 November 2021

