## Supplementary Online Material for "Are Pacific Chorus Frogs (*Pseudacris regilla*) Resistant to Tetrodotoxin (TTX)? Characterizing Potential TTX Exposure and Resistance in an Ecological Associate of Pacific Newts (*Taricha*)"

*Supporting Methods.*—DNA Extraction Protocol, adapted from Miller et al. (1988)

Day One Procedure

1. Cut a small piece (approximately 2 mm in diameter) of tissue for extraction.
2. Create master mix by multiplying each reagent by number of samples plus 2 µL for extra potentially needed due to pipetting error):
    - 410 µL extraction buffer
    - 80 µL SDS
    - 10 µL proteinase K (20 g/L)
    - 2 µL RNase A
3. Digest the small piece of each tissue in 502 L of master mix overnight at 55 ℃.

Day Two Procedure

1. Spin samples at 13,000 rpm for 5 min.
2. Pour supernatant from each into new 1.5 mL tube.
3. Add 180 µL of 5 M NaCl to each tube and invert 50 times each.
4. Spin tubes at 13,000 rpm for 5 min.
5. Add 420 µL of cooled isopropanol to each new 1.5 mL tube.
6. Pour supernatant into a tube with isopropanol and invert gently 10-15 times.
7. Spin samples at 13,000 rpm for 7 min.
8. Discard supernatant carefully into ethanol waste.
9. Add 250 µL of 80% EtOH to each tube, invert 50x, and vortex.
10. Spin samples at 13,000 rpm for 7 min.
11. Discard supernatant into EtOH waste carefully.

12. Add 250 µL of 80% EtOH to each sample, invert 50x, and vortex.
13. Spin samples at 13,000 rpm for 7 min.
14. Pour off supernatant into EtOH waste once more and dab with paper towels.
15. Put uncapped tubes in a speed vacuum (no heat) for about 30 min, until the tube is dry.
16. Add 50 µL TE to each tube and vortex.
17. Place samples in the incubator at 55 ℃ for about 3 h, and then transfer samples to new 1.5 mL locking tubes.

Table S1.— Metadata for tissues obtained from the Museum of Vertebrate Zoology (UC Berkeley) and from newly collected specimens (indicated by asterisks) that were used to sequence SCN4A exon 24 in this study. Overlap implies that the coordinates were within 500 m elevation or 35 m of a documented *Taricha* spp. occurrence in the Global Biodiversity Information Facility database.

| Species | County (field site, if applicable) | Latitude | Longitude | Overlap with *Taricha*? | GenBank number | MVZ collection number |
|---|---|---|---|---|---|---|
| *P. regilla* | Alpine | 38.49078 | -119.80266 | No | OL988632 | MVZ:Herp:137395 |
| *P. regilla* | Alpine | 38.49078 | -119.80266 | No | OL988633 | MVZ:Herp:137396 |
| *P. regilla* | Contra Costa (OBRT 2)* | 37.94441 | -122.13377 | Yes | OL988634 | MVZ:Herp:301109 |
| *P. regilla* | Humboldt | 40.69283 | -124.27447 | Yes | OL988635 | MVZ:Herp:272809 |
| *P. regilla* | Inyo | 35.936948 | -117.90531 | No | OL988636 | MVZ:Herp:145343 |

45 Table S1, continued

| | | | | | | |
|---|---|---|---|---|---|---|
| *P. regilla* | Inyo | 36.112461 | -117.17456 | No | OL988637 | MVZ:Herp:145412 |
| *P. regilla* | Inyo | 36.112461 | -117.17456 | No | OL988638 | MVZ:Herp:173518 |
| *P. regilla* | Los Angeles | 34.11175 | -118.77242 | Yes | OL988639 | MVZ:Herp:233336 |
| *P. regilla* | Los Angeles | 34.11175 | -118.77242 | Yes | OL988640 | MVZ:Herp:233337 |
| *P. regilla* | Mariposa | 37.66312 | -119.60102 | No | OL988641 | MVZ:Herp:240750 |
| *P. regilla* | Mariposa | 37.74597 | -119.7993 | No | OL988642 | MVZ:Herp:240768 |
| *P. regilla* | Mariposa | 37.67561 | -119.65156 | No | OL988643 | MVZ:Herp:240775 |
| *P. regilla* | Mariposa | 37.72424 | -119.63531 | No | OL988644 | MVZ:Herp:249951 |
| *P. regilla* | Mariposa | 37.70265 | -119.75067 | No | OL988645 | MVZ:Herp:249953 |
| *P. regilla* | Mono | 38.18458 | -119.58434 | No | OL988646 | MVZ:Herp:249958 |

Table S1, continued

| | | | | | | |
|---|---|---|---|---|---|---|
| *P. regilla* | Mono | 38.11097 | -119.44518 | No | OL988647 | MVZ:Herp:249961 |
| *P. regilla* | Mono | 38.11097 | -119.44518 | No | OL988648 | MVZ:Herp:249962 |
| *P. regilla* | Monterey | 36.8392685 | -121.78979 | Yes | OL988649 | MVZ:Herp:145419 |
| *P. regilla* | Monterey | 36.8392685 | -121.78979 | Yes | OL988650 | MVZ:Herp:145420 |
| *P. regilla* | Monterey | 36.8392685 | -121.78979 | Yes | OL988651 | MVZ:Herp:145421 |
| *P. regilla* | San Diego | 32.97 | -116.73 | Yes | OL988652 | MVZ:Herp:220013 |
| *P. regilla* | San Francisco (Presidio)* | 37.788496 | -122.468605 | No | OL988653 | MVZ:Herp:301135 |
| *P. regilla* | San Francisco (Presidio)* | 37.788496 | -122.468605 | No | OL988654 | MVZ:Herp:301137 |

Table S1, continued

| | | | | | | |
|---|---|---|---|---|---|---|
| *P. regilla* | San Francisco (Presidio)* | 37.788496 | -122.468605 | No | OL988655 | MVZ:Herp:301139 |
| *P. regilla* | San Francisco (Presidio)* | 37.788496 | -122.468605 | No | OL988656 | MVZ:Herp:301141 |
| *P. regilla* | San Francisco (Presidio)* | 37.788496 | -122.468605 | No | OL988657 | MVZ:Herp:301143 |
| *P. regilla* | San Francisco (Presidio)* | 37.788496 | -122.468605 | No | OL988658 | MVZ:Herp:301146 |
| *P. regilla* | San Francisco (Presidio)* | 37.788496 | -122.468605 | No | OL988659 | MVZ:Herp:301147 |
| *P. regilla* | San Francisco (Presidio)* | 37.788496 | -122.468605 | No | OL988660 | MVZ:Herp:301149 |

Table S1, continued

| | | | | | | |
|---|---|---|---|---|---|---|
| *P. regilla* | San Luis Obispo | 35.424791 | -120.56755 | Yes | OL988661 | MVZ:Herp:150009 |
| *P. regilla* | Santa Barbara | 34.6109479 | -120.1992 | No | OL988662 | MVZ:Herp:137355 |
| *P. regilla* | Santa Barbara | 34.6109479 | -120.1992 | No | OL988663 | MVZ:Herp:137356 |
| *P. regilla* | Santa Barbara | 34.6109479 | -120.1992 | No | OL988664 | MVZ:Herp:137359 |
| *P. regilla* | Sutter | 39.20568 | -121.81982 | No | OL988665 | MVZ:Herp:229175 |
| *P. regilla* | Tuolumne | 37.90399 | -119.82941 | Yes | OL988666 | MVZ:Herp:240791 |
| *P. regilla* | Tuolumne | 37.91745 | -119.8076 | Yes | OL988667 | MVZ:Herp:240796 |
| *P. regilla* | Tuolumne | 38.01339 | -119.72839 | Yes | OL988668 | MVZ:Herp:250044 |
| *P. regilla* | Tuolumne | 38.17303 | -119.59591 | No | OL988669 | MVZ:Herp:250047 |

Table S1, continued

| | | | | | | |
|---|---|---|---|---|---|---|
| *Acris crepitans* | Callaway County, MO | 38.58193 | -92.10184 | No but overlaps with TTX-bearing *N. viridescens* | OM069736 | MVZ:Herp:240049 |
| *Pseudacris cadaverina* | Cataviña, Baja CA | 29.735556 | -114.7016667 | No | OM069735 | MVZ:Herp:145365 |

*Table S2.*—Sequences of $Na_V 1.4$ (SCN4A exon 24) obtained from GenBank used in Fig. 3.

| | |
|---|---|
| *Adelphobates galactonotus* | KT989158.1 |
| *Afronatrix anoscopus* | JQ687831.1 |
| *Agkistrodon contortrix* | JQ687777.1 |
| *Allobates femoralis* | KT989148.1 |
| *Allobates talamancae* | KT989149.1 |
| *Allobates zaparo* | KT989150.1 |
| *Ambystoma tigrinum mavortium* | KP118977.1 |
| *Ameerega bilinguis* | KT989151.1 |
| *Ameerega hahneli* | KT989152.1 |
| *Ameerega petersi* | MK278829.1 |
| *Ameerega picta* | MK278828.1 |
| *Ameerega trivittata* | KT989154.1 |
| *Ameerega trivittata* | MK278858.1 |
| *Amphiesma pryeri* | JQ687834.1 |
| *Amphiesma pryeri* | JQ687835.1 |
| *Amphiesma sp.* | JQ687833.1 |
| *Amphiesma vibakari* | JQ687832.1 |
| *Andinobates bombetes* | MK278802.1 |
| *Andinobates bombetes* | MK278832.1 |
| *Andinobates fulguritus* | MK278827.1 |

Table S2, continued

| | |
|---|---|
| *Anolis carolinensis* | XM_8113208.2 |
| *Aromobates saltuensis* | KT989147.1 |
| *Bolitoglossa vallecula* | GHME01165342.1 |
| *Bufo bufo* | XM_40435981.1 |
| *Bufo gargarizans* | XM_44297786.1 |
| *Causus maculatus* | JQ687780.1 |
| *Charina bottae* | JQ687776.1 |
| *Clonophis kirtlandii* | JQ687851.1 |
| *Colostethus fugax* | KT989163.1 |
| *Colostethus panamansis* | KT989164.1 |
| *Coluber constrictor* | JQ687813.1 |
| *Coniophanes bipunctatus* | JQ687788.1 |
| *Coniophanes fissidens* | JQ687789.1 |
| *Crotalus oreganus* | JQ687778.1 |
| *Crotalus tigris* | XM_39331422.1 |
| *Cryptelytrops albolabris* | JQ687779.1 |
| *Cynops pyrrhogaster* | KP118971.1 |
| *Dendrelaphis* sp. | JQ687816.1 |
| *Dendrobates auratus* | MK278838.1 |
| *Dendrobates tinctorius* | KT989160.1 |
| *Dendrobates tinctorius* | MZ545382.1 |

Table S2, continued

| | |
|---|---|
| *Dendrobates truncatus* | MK278843.1 |
| *Drymarchon corais* | JQ687814.1 |
| *Drymobius margaritiferus* | JQ687809.1 |
| *Elapsoidea nigra* | JQ687782.1 |
| *Eleutherodactylus johnstonei* | MH050340.1 |
| *Elgaria multicarinata* | JQ687775.1 |
| *Enhydris* sp. | JQ687781.1 |
| *Epipedobates anthonyi* | KT989166.1 |
| *Epipedobates boulengeri* | MK278805.1 |
| *Epipedobates machalilla* | KT989169.1 |
| *Espadarana callistomma* | KT989144.1 |
| *Excidobates captivus* | KT989157.1 |
| *Gastrotheca litonedis* | KT989143.1 |
| *Gekko japonicus* | XM_15418240.1 |
| *Gonionotophis klingi* | JQ687783.1 |
| *Grayia smythii* | JQ687818.1 |
| *Hapsidophrys lineatus* | JQ687803.1 |
| *Helicops angulatus* | JQ687801.1 |
| *Heterodon nasicus* | JQ687784.1 |
| *Heterodon platirhinos* | JQ687785.1 |
| *Heterodon platirhinos* | KT277703.1 |

Table S2, continued

| | |
|---|---|
| *Hyloxalus italoi* | KT989155.1 |
| *Hyloxalus nexipus* | KT989156.1 |
| *Hypsiboas picturatus* | KT989146.1 |
| *Incilius nebulifer* | KT989142.1 |
| *Lacerta agilis* | XM_33169347.1 |
| *Lacerta agilis* | XM_33169506.1 |
| *Liopeltis tricolor* | JQ687802.1 |
| *Liophis epinephelus* | JQ687790.1 |
| *Liophis miliaris* | JQ687793.1 |
| *Liophis poecilogyrus* | JQ687794.1 |
| *Liophis typhlus* | JQ687792.1 |
| *Lithodytes lineatus* | KT989145.1 |
| *Lygophis anomalus* | JQ687791.1 |
| *Lystrophis dorbignyi* | JQ687796.1 |
| *Lystrophis semicinctus* | JQ687795.1 |
| *Mantella aurantiaca* | KT989141.1 |
| *Nanorana parkeri* | XM_18560831.1 |
| *Natrix natrix* | JQ687836.1 |
| *Natrix natrix* | JQ687840.1 |
| *Natrix tessellata* | JQ687844.1 |
| *Notechis scutatus* | XM_26694172.1 |

Table S2, continued

| | |
|---|---|
| *Notophthalmus viridescens* | KP118970.1 |
| *Oophaga histrionica* | MK278814.1 |
| *Oophaga histrionica* | MK278815.1 |
| *Oophaga pumilio* | KT989159.1 |
| *Oophaga pumilio* | MK278841.1 |
| *Ophiophagus hannah* | BK009415.1 |
| *Pachytriton labiatus* | KP118972.1 |
| *Pantherophis guttatus* | XM_34416072.1 |
| *Phyllobates aurotaenia* | KT989161.1 |
| *Phyllobates aurotaenia* | MK278808.1 |
| *Phyllobates aurotaenia* | MK278820.1 |
| *Phyllobates aurotaenia* | MK278826.1 |
| *Phyllobates bicolor* | MK278728.1 |
| *Phyllobates bicolor* | MK278803.1 |
| *Phyllobates lugubris* | MK278810.1 |
| *Phyllobates lugubris* | MK278839.1 |
| *Phyllobates lugubris* | MK278840.1 |
| *Phyllobates terribilis* | KT989162.1 |
| *Phyllobates terribilis* | MK278855.1 |
| *Phyllobates terribilis* | MZ545381.1 |
| *Pituophis catenifer* | JQ687806.1 |

Table S2, continued

| | |
|---|---|
| *Pleurodeles waltl* | KP118974.1 |
| *Podarcis muralis* | XM_28702752.1 |
| *Pogona vitticeps* | XM_20798034.1 |
| *Protobothrops mucrosquamatus* | XM_29287219.1 |
| *Ptyas korros* | JQ687808.1 |
| *Ptyas mucosus* | JQ687807.1 |
| *Python bivittatus* | XM_25164533.1 |
| *Ramphotyphlops bituberculatus* | KX079442.1 |
| *Rana temporaria* | XM_40331478.1 |
| *Ranitomeya toraro* | MK278835.1 |
| *Ranitomeya ventrimaculata* | MK278836.1 |
| *Rhabdophis himalayanus* | JQ687819.1 |
| *Rhabdophis subminiatus* | JQ687829.1 |
| *Rhabdophis tigrinus* | JQ687820.1 |
| *Rhabdophis tigrinus* | JQ687826.1 |
| *Rhabdophis tigrinus* | JQ687827.1 |
| *Rheobates palmatus* | MK278846.1 |
| *Rheobates palmatus* | MK278847.1 |
| *Rhinatrema bivittatum* | XM_29572321.1 |
| *Salamandra salamandra* | KP118976.1 |
| *Sceloporus undulatus* | XM_42471834.1 |

Table S2, continued

| | |
|---|---|
| *Silverstoneia flotator* | KT989165.1 |
| *Silverstoneia nubicola* | MK278842.1 |
| *Silverstoneia* sp. | MK278844.1 |
| *Silverstoneia* sp. | MK278845.1 |
| *Sinonatrix aequifasciata* | JQ687849.1 |
| *Taricha granulosa* | KP118969.1 |
| *Taricha torosa* | KP118968.1 |
| *Thamnodynastes strigatus* | JQ687799.1 |
| *Thamnophis atratus* | FJ570810.1 |
| *Thamnophis atratus* | FJ571014.1 |
| *Thamnophis couchii* | FJ570812.1 |
| *Thamnophis couchii* | MT304461.1 |
| *Thamnophis elegans* | FJ570811.1 |
| *Thamnophis elegans* | FJ571033.1 |
| *Thamnophis elegans* | XM_32238131.1 |
| *Thamnophis eques* | JQ687858.1 |
| *Thamnophis errans* | FJ571032.1 |
| *Thamnophis fulvus* | JQ687857.1 |
| *Thamnophis proximus* | FJ571045.1 |
| *Thamnophis radix* | FJ571046.1 |
| *Thamnophis sirtalis Benton* | AY851744.1 |

Table S2, continued

| | |
|---|---|
| *Thamnophis sirtalis Warrenton* | AY851745.1 |
| *Thamnophis sirtalis Willow Creek* | AY851746.1 |
| *Thamnophis sirtalis* | KY745652.1 |
| *Thamnophis sirtalis* | KY745662.1 |
| *Thamnophis sirtalis* | KY745680.1 |
| *Triturus dobrogicus* | KP118973.1 |
| *Tylototriton shanjing* | KP118975.1 |
| *Tylototriton wenxianensis* | GESS01000732.1 |
| *Tylototriton wenxianensis* | GESS01029581.1 |
| *Varanus komodoensis* | XM_44418683.1 |
| *Virginia striatula* | FJ571064.1 |
| *Xenochrophis piscator* | JQ687830.1 |
| *Xenodon rabdocephalus* | JQ687797.1 |
| *Xenodon rabdocephalus* | JQ687798.1 |
| *Xenopus laevis* | XM_41576824.1 |
| *Xenopus laevis* | XM_41578552.1 |
| *Xenopus tropicalis* | XM_018089322.2 |
| *Zootoca vivipara* | XM_35134760.1 |

Figure S1.—Field Sites: A. Old Briones Road Trail 1 (OBRT 1) pond, 37.94441, -122.13377; B. Old Briones Road Trail 2 (OBRT 2) pond, 37.94352, -122.14102; C. Japanese Pool at the UC Botanical Garden (UCBG), 37.87440, -122.23760; D. Presidio site, 37.788496, -122.468605; all photos taken by K. Montana.

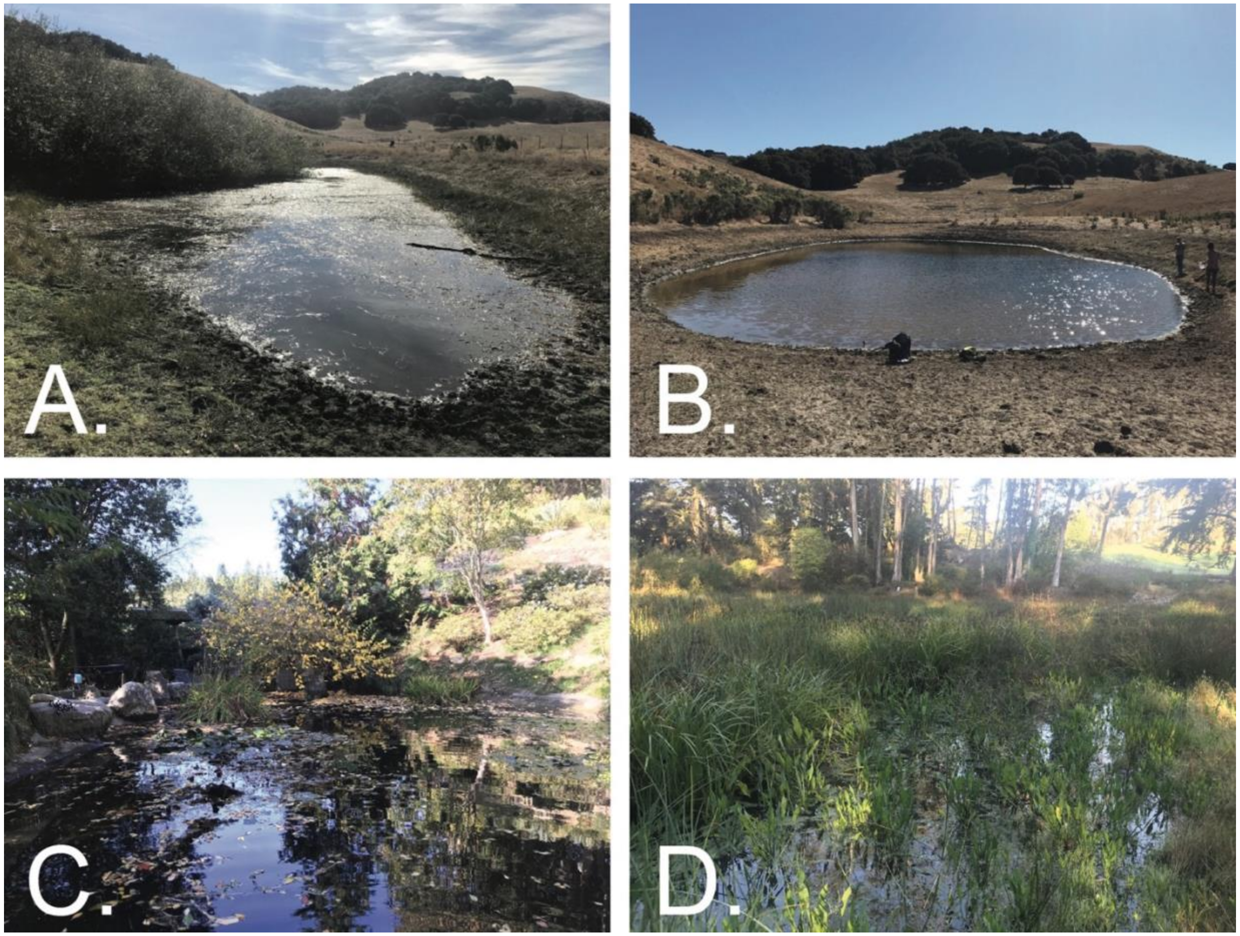

*Figure S2.*—Numbers of A. *P. regilla* adults, B. *Taricha torosa* and *T. granulosa* adults and larvae, and C. *T. torosa* egg masses observed over field trips to the Old Briones Road Trail 1 (OBRT 1) (blue circle line), OBRT 2 (red triangle line), and U.C. Botanical Garden (yellow square line) ponds; 20 October 2019 to late February 2020.

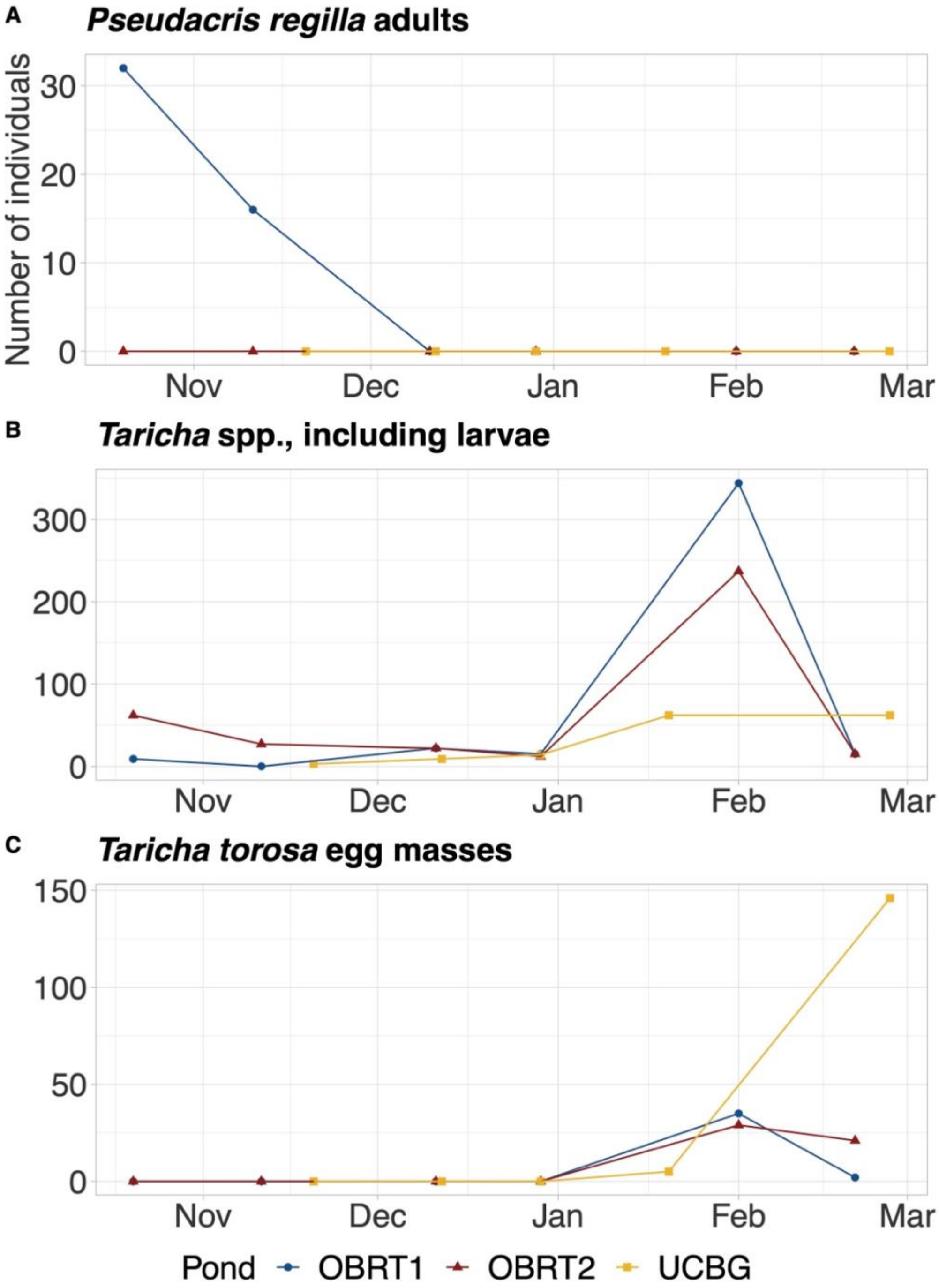

*Figure S3*.—Percentage of total abundance of GBIF-recorded individuals of *Pseudacris regilla* (green) and *Taricha* (blue) in California by month, over the course of the last 50 years. Peak abundances for *P. regilla* occur around April and August each year, and peak abundances for *Taricha* occur in January each year. Samples included were from 1 January 1971 to 23 November 2021.

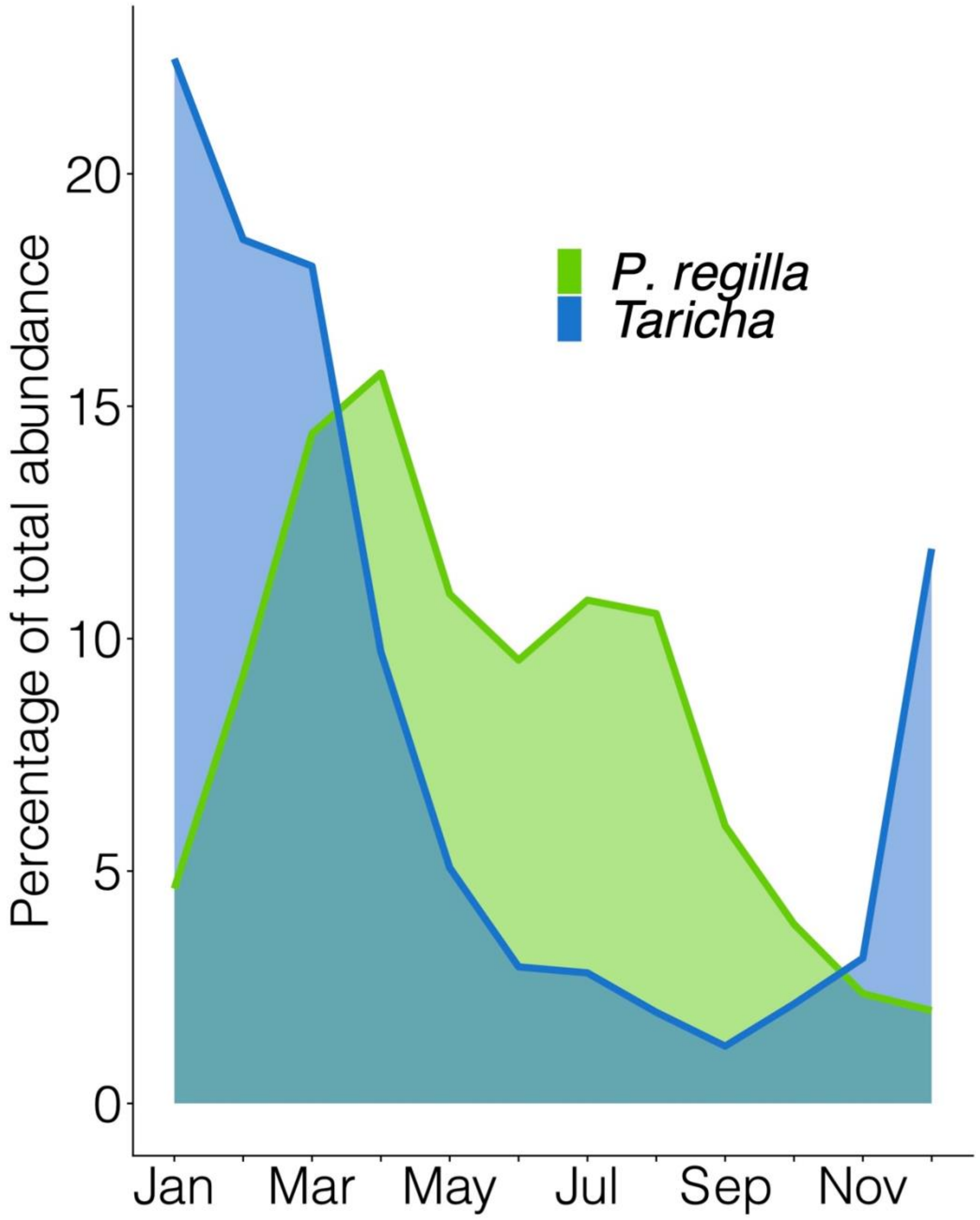

*Figure S4*.—Percentage of total abundance of GBIF-recorded individuals of *Pseudacris regilla* (green) and *Taricha* (blue) in the San Francisco Bay Area (N 38.66226, S 37.09298, E -121.12701, W -123.23579) by month, over the course of the last 50 years. Peak abundance for *P. regilla* occurs around March each year, and peak abundance for *Taricha* occurs in January each year. Samples included were from 1 January 1971 to 23 November 2021.

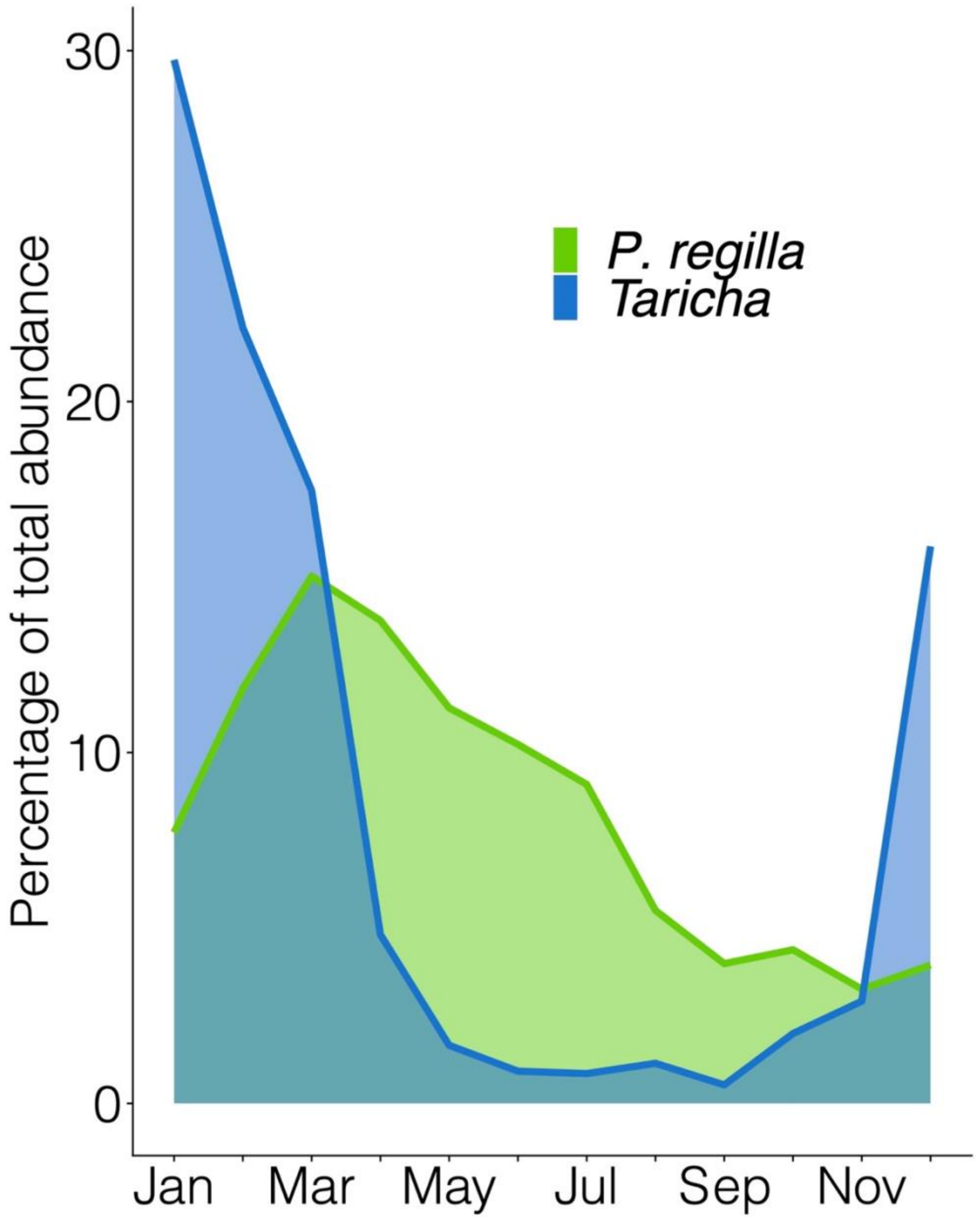
