## Supplementary figures and images for "Are Pacific Chorus Frogs (*Pseudacris regilla*) Resistant to Tetrodotoxin (TTX)? Characterizing Potential TTX Exposure and Resistance in an Ecological Associate of Pacific Newts (*Taricha*)"

### Figure S1

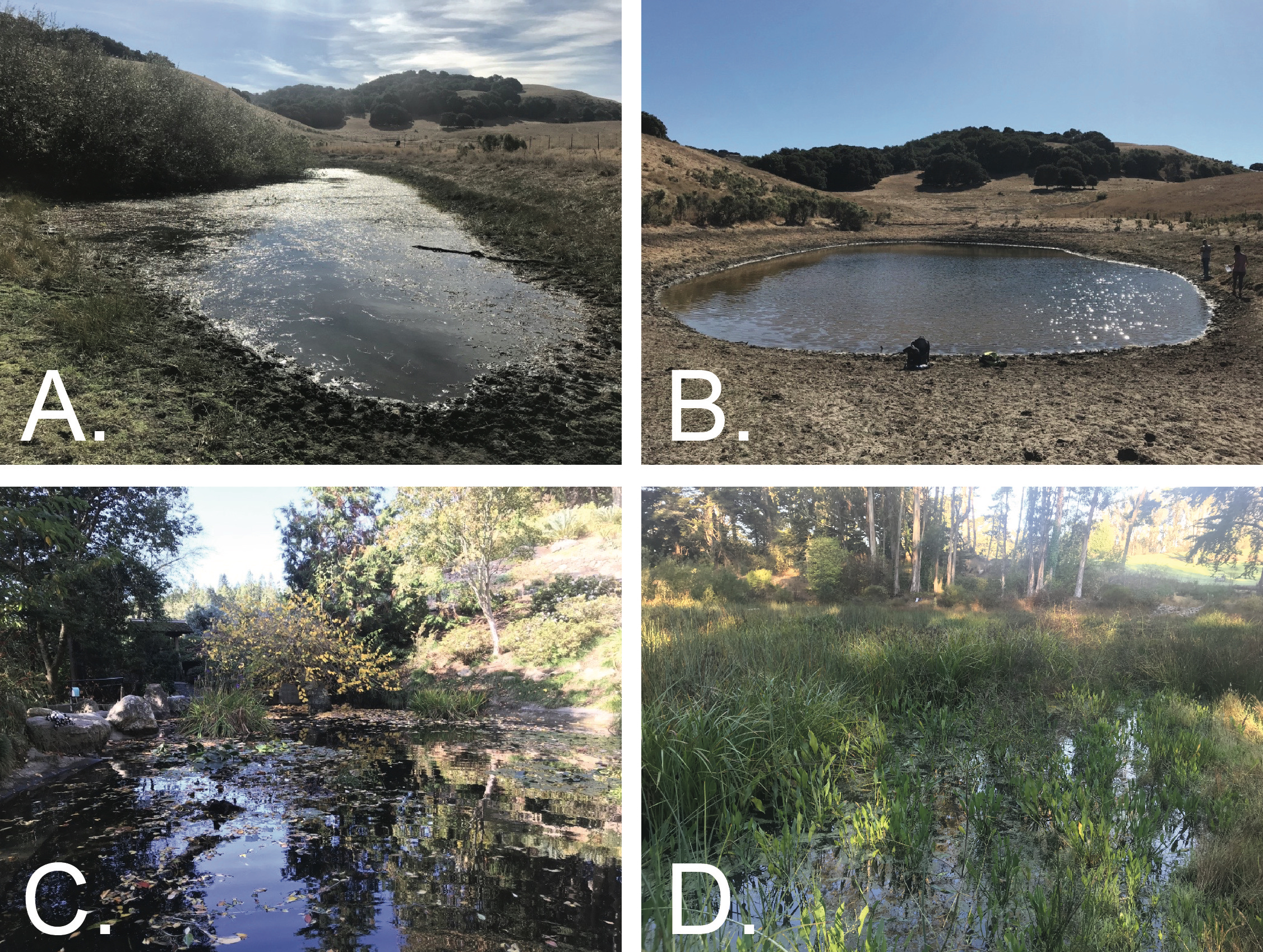

### Figure S2

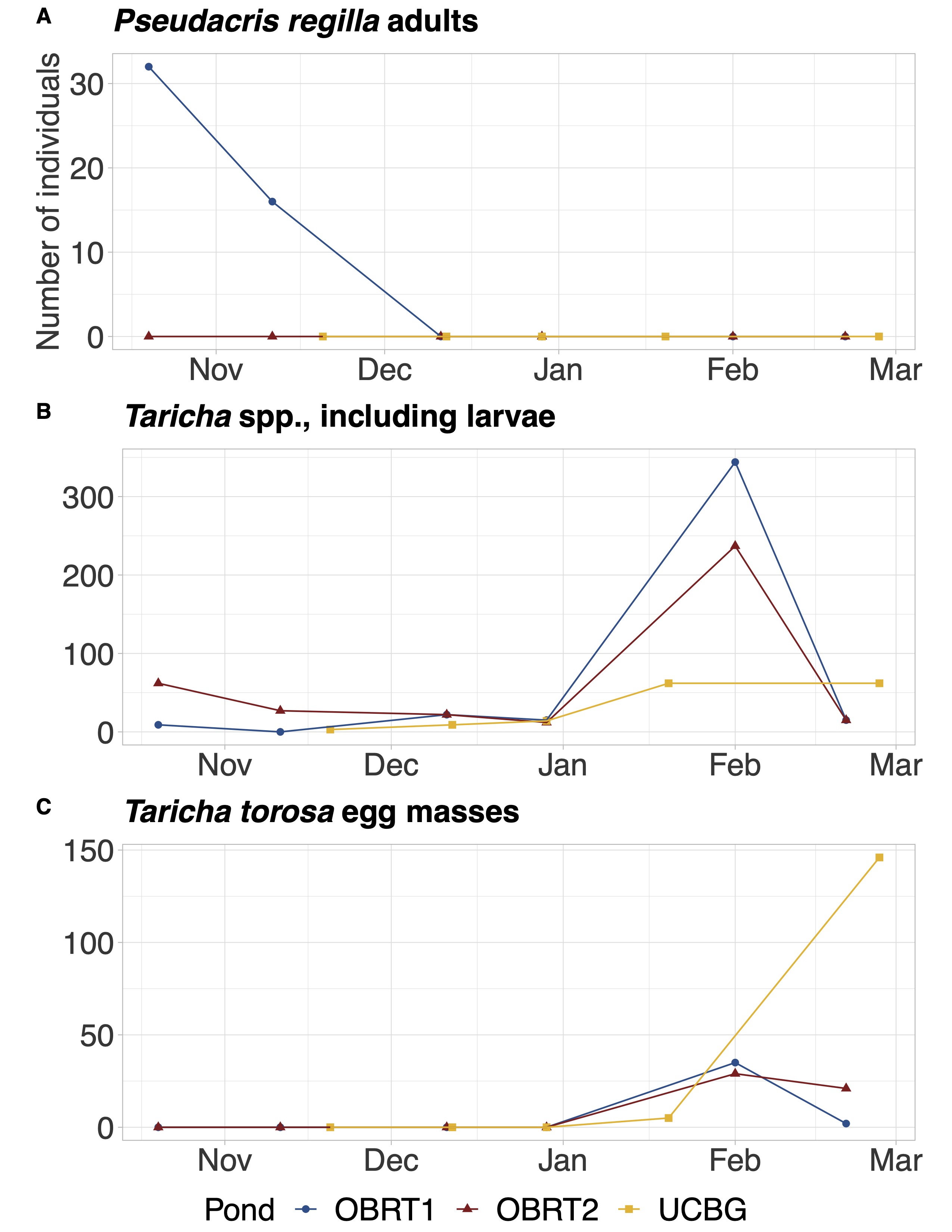

### Figure S4

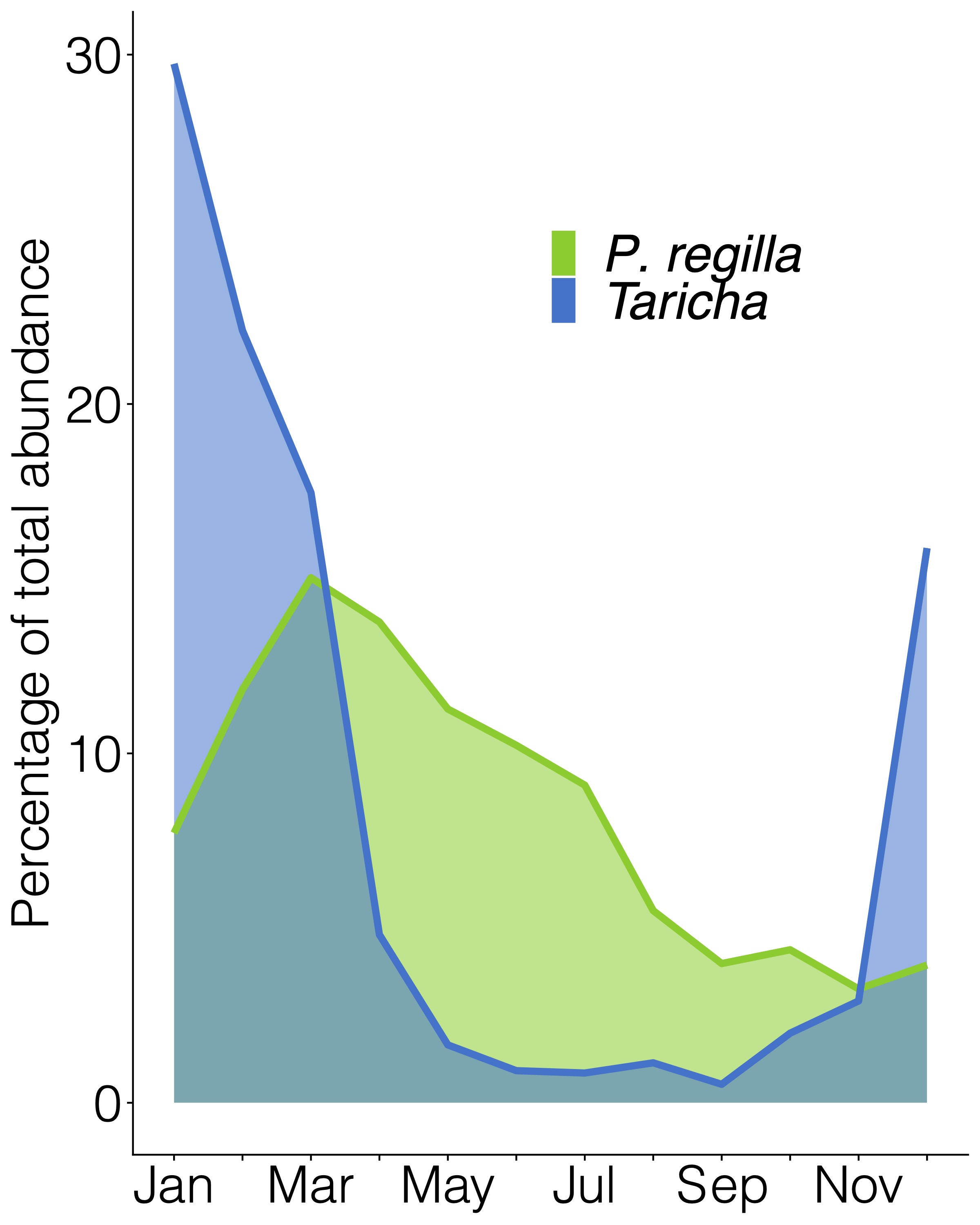

### Figures S3

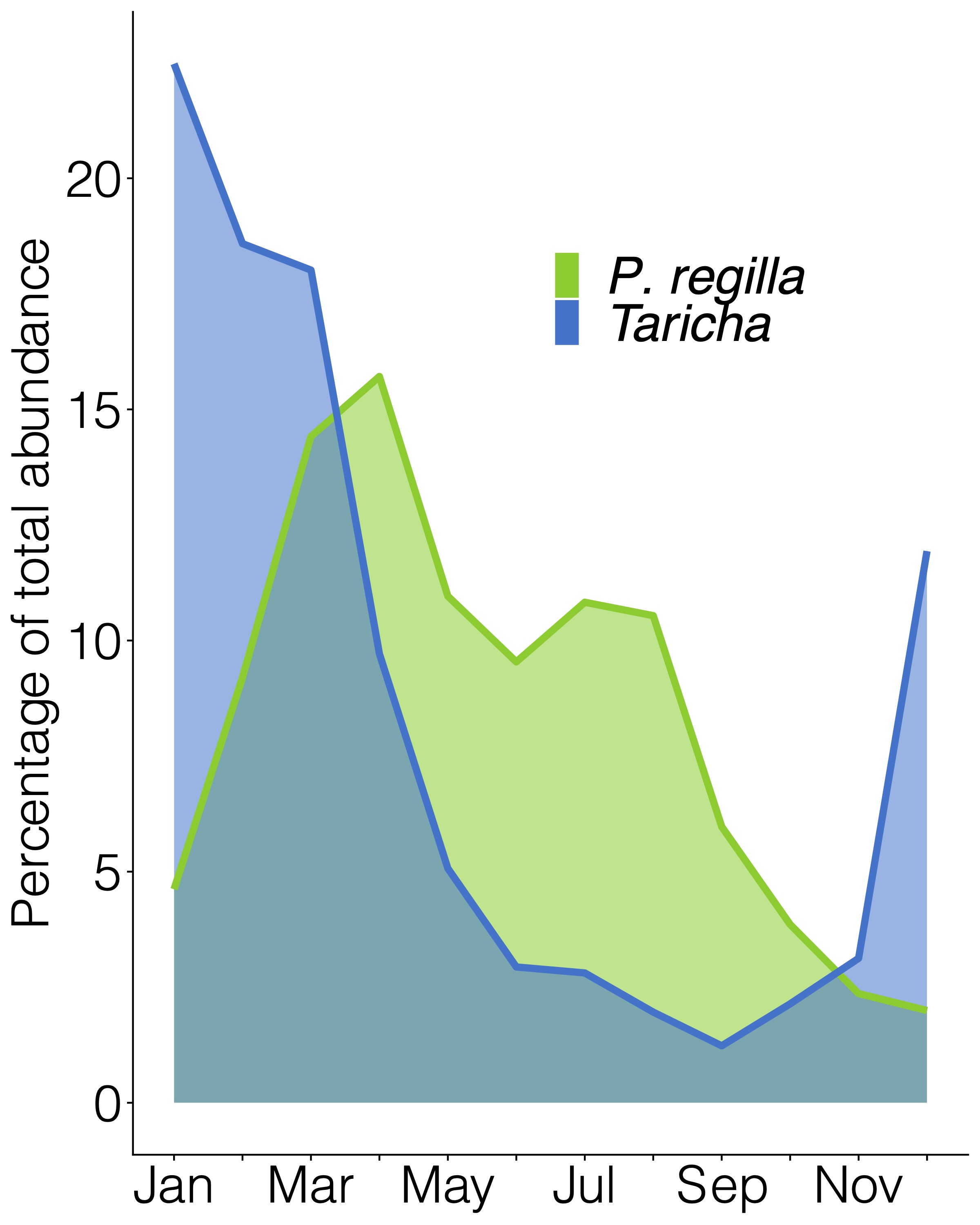
